# Endothelial Heterogeneity Across Vascular Beds Impacts Inflammatory Signaling and Neutrophil Adhesion

**DOI:** 10.64898/2026.05.26.727909

**Authors:** Eric L Ginter, Sunanda Mitra, Laurel E Hind

## Abstract

Endothelial cells (ECs) are key players in maintaining homeostasis and coordinating immune responses, activating during acute inflammation to recruit immune cells. Endothelial heterogeneity has been found to impact transcription level differences across EC sources, but how these differences drive downstream effects in inflammatory signaling and immune interactions remains unclear. Here, we employed multiplexed ELISA to quantify secretion for 19 inflammatory factors following tumor necrosis factor (TNF) or *Pseudomonas aeruginosa* activation of four primary human EC sources: umbilical artery (HUAEC), umbilical vein (HUVEC), dermal microvascular (HDMEC), and pulmonary microvascular (HPMEC) endothelial cells. We also quantified changes in neutrophil adhesion to each EC source and used partial least squares regression (PLSR) to identify key inflammatory proteins associated with changes in neutrophil adhesion. We found distinct inflammatory secretion profiles across all cell types, with veinous ECs showing the highest basal secretion of most inflammatory proteins and pulmonary ECs exhibiting the lowest. Arterial ECs exhibited the lowest sensitivity to inflammatory stimulus, while pulmonary ECs exhibited dynamic responses following activation. Furthermore, inflammatory stimulus caused large differences in expression across cell sources for six factors: GM-CSF, IL-1β, IL-6, IP-10, E-selectin, and ICAM-1. We found endothelial heterogeneity also contributed to differences in neutrophil adhesion to unstimulated ECs. Our PLSR analysis revealed five secreted factors most indicative of changes in neutrophil adhesion: E-selectin, ICAM-1, PECAM1, IL-6, and IL-8. Collectively, our findings strengthen the emerging view that vascular-bed specific differences in EC phenotype can impact downstream immune responses.

## Introduction

Endothelial cells (ECs) form the inner lining of blood vessels and are essential regulators of vascular homeostasis. Though traditionally viewed as a passive barrier separating circulating blood from surrounding tissues, the endothelium is now recognized as an active participant in many physiological processes, including vascular permeability, coagulation, angiogenesis, and immune regulation [1]. During inflammation, endothelial cells upregulate secreted factors and adhesion molecules on their surface [2]. Neutrophils, the most numerous white blood cells in the circulation, respond to these signals by changing their surface protein expression and adhere to the endothelium through rolling and tight binding interactions [2]. These endothelial-neutrophil interactions are key to activating neutrophils to properly fight infection. Evidence suggests that endothelial cells exhibit broad phenotypic heterogeneity based on tissue source [3–5], but these differences are not well characterized in the context of the inflammatory response and resultant neutrophil recruitment. This study aims to quantify the differences in inflammatory protein secretion due to endothelial heterogeneity resulting from tissue sourcing.

Although endothelial cells share core functions, increasing evidence shows substantial phenotypic heterogeneity across vascular beds. Endothelial cell heterogeneity has long been recognized, with differences attributed to vessel characteristics including size (large vessel vs microvasculature), vessel type (artery vs vein) and tissue origin [3]. Single-cell resolution techniques have brought further understanding of endothelial cell phenotypes, indicating transcription-level differences of myriad proteins responsible for angiogenesis, proliferation, inflammatory responses, and more [4–7]. For example, iPSC-derived endothelial cells with arterial and venous characteristics had different levels of transcripts for cytokines MCP-1 and IP10 [6]. While many studies have measured the transcriptome to describe endothelial heterogeneity, few have looked at downstream outcomes including inflammatory protein secretion and immune cell interactions.

Despite increased recognition of endothelial heterogeneity, many in vitro studies continue to rely heavily on a limited number of endothelial models, particularly human umbilical vein endothelial cells (HUVECs), to investigate vascular inflammation and immune interactions. However, reliance on a single endothelial source may overlook important vascular-bed specific responses. Whether endothelial origin alters inflammatory signaling and subsequent immune cell interactions remains insufficiently characterized. Relatively few studies have directly compared inflammatory responses across endothelial cells derived from multiple vascular sources, and even fewer have examined how these differences influence downstream immune outcomes. Therefore, we sought to determine how endothelial source influences inflammatory secretion and resultant neutrophil behavior.

The objective of the present study was to determine how endothelial cell source influences inflammatory responses and downstream neutrophil interactions across distinct vascular beds. Human endothelial cells derived from umbilical artery, umbilical vein, dermal microvascular, and pulmonary microvascular sources were examined to characterize both basal and inflammatory phenotypes. Endothelial activation was induced using either TNF or *Pseudomonas aeruginosa* exposure to compare responses to cytokine-mediated and pathogen-associated inflammatory stimuli. We also developed an assay to quantify neutrophil adhesion to the endothelial surface and characterize differences across endothelial cell sources. Finally, we applied partial least squares regression analysis to identify endothelial cell-secreted inflammatory factors associated with neutrophil recruitment. We found each endothelial cell source displayed a unique secretome with large variation in protein expression following inflammatory activation. Venous endothelial cells exhibited the highest basal secretion for most inflammatory factors measured and had the highest levels of neutrophil adhesion when unstimulated. Pulmonary endothelial cells demonstrated the lowest basal secretion for most factors but also saw the largest relative increases of all cell types for critical inflammatory markers including GM-CSF, IL-1β, and ICAM-1. The regression model found three proteins whose expression varied most across cell source were also highly correlated with neutrophil adhesion changes: E-selectin, ICAM-1, and IL-6. Together, our findings indicate diverse basal and inflammatory secretion profiles across endothelial cell sources that influence downstream neutrophil recruitment dynamics, and underscore how critical endothelial source is as a design consideration for *in vitro* models.

## Methods

### Endothelial cell culture

The following cells were used: pooled human umbilical vein endothelial cells (HUVEC, PromoCell C-12203, Heidelberg, Germany), human umbilical arterial endothelial cells (HUAEC, PromoCell C-12202, Heidelberg, Germany), adult dermal microvascular endothelial cells (HDMEC, PromoCell C-12212, Heidelberg, Germany), and adult pulmonary microvascular endothelial cells (HPMEC, PromoCell C-12281, Heidelberg, Germany). HUVECs and HUAECs were cultured in Endothelial Growth Medium 2 (EGM-2, Lonza Walkersville CC-3162, Basel, Switzerland) while HDMEC and HPMEC were cultured in Endothelial Growth Medium 2 Microvascular (EGM-2 MV, Lonza Walkersville CC-4147, Basel, Switzerland). All cells were grown to ~80% confluence and grown through passage 6. All experiments utilized endothelial cells from passages 3-6.

### TNF activation

Tumor necrosis factor (TNF, R&D systems 210-TA-020/CF, Minneapolis, Minnesota) was diluted in phosphate-buffered saline (PBS, B2944-100, Thermo Scientific, Waltham, MA) to 100 mg/mL and stored at −20 °C. For experiments, solutions of 50 ng/mL in EGM-2 or EGM-2 MV were created and added to each well.

### *Pseudomonas aeruginosa* culture

*P. aeruginosa* was prepared as previously described [8]. Briefly, *P. aeruginosa* strain K (PAK) was streaked onto LB plates and cultured overnight at 37 °C. A single bacteria colony was sub-cultured overnight in LB broth at 37 °C. The subculture was then diluted 1:4 with fresh broth and grown in an incubator for 1.5 hours at 37 °C. Once expanded, bacteria were pelleted by centrifugation for 1 minute at 17,000xg, and resuspended in EGM-2 or EGM-2 MV to an optical density (OD) of 0.05 (measured at 600 nm, 1.25 × 10^6^ CFU mL−1) for use in experiments.

### MagPix secretion analysis

Endothelial cells were cultured as described above and seeded in 96 well plates (5E5 cells/well) and grown to confluence for 3 days. Each well was activated with 50 μL of either corresponding culture media with no additives, TNF (50 ng/mL in media), or PAK (OD 0.05 in media) and incubated at 37 °C for 16 hours. Media was collected and centrifuged to remove cellular debris. Samples were stored at −80°C until the assay was run. A MagPix Luminex Xmap system (Thermo Fisher Scientific, Waltham, MA) was used with a custom Procartaplex panel of inflammatory and endothelial injury markers (Invitrogen PPX-21, Waltham, MA) following the manufacturer’s protocol. This panel measured the following markers: angiopoietin-1, E-selectin, GM-CSF, ICAM-1, IFN-α, IFN-γ, IL-1α, IL-1β, IL-6, IL-8, IL-10, IL-12p70, IP-10, MCP-1, MIP-1α, MIP-1β, PECAM-1, P-selectin, and syndecan. Data shown was collected from 4 replicates per condition.

### Neutrophil isolation from whole blood

All blood samples were obtained according to the institutional review board-approved protocols per the Declaration of Helsinki. Peripheral blood neutrophils were isolated from healthy adult donors, using a MACSxpress Neutrophil Isolation Kit (Miltenyi Biotec 130–104-434, Bergisch Gladbach, Germany) and a MACSxpress Erythrocyte Depletion Kit (130–098-196, Miltenyi Biotec, Bergisch Gladbach, Germany), according to the manufacturer’s protocols. Informed consent was obtained from donors at the time of the blood draw according to our institutional review board protocol number 20-0082.

#### Neutrophil Adhesion Assay

Endothelial cells were cultured as described above, seeded in 96 well plates (5E5 cells/well), and grown to confluence for 3 days. Each well was activated with 50 μL of either corresponding media with no additives, TNF (50 ng/mL in media), or PAK (OD 0.05 in media) and incubated at 37 °C for 2 hours. Each activating solution was supplemented with calcein red-orange (C34851 Thermo Scientific, Waltham, MA) diluted 1:2000. Neutrophils were isolated as described above and stained with calcein AM (C3100MP, Thermo Scientific, Waltham, MA) diluted 1:1000 in PBS (B2944-100, Thermo Scientific, Waltham, MA) for 10 minutes at RT. Stained neutrophils were then added to the stained and activated endothelial cells (4E5 cells/well) and incubated at 37 °C for 10 minutes to adhere. Each well was imaged with a confocal microscope (scope details). Non-adherent neutrophils were then removed with 5x washes of DPBS(+Ca/+Mg) (14040-117, Thermo Scientific, Waltham, MA) and imaging was repeated.

All images were taken using a Nikon A1R HD25 Laser Scanning Confocal Microscope built on the Nikon TI2-E Inverted Microscope System, a fully automated stage, and Nikon Elements acquisition software. Imaging was performed with a Nikon 20x/0.75 (NA) objective in the 561nm and 488nm wavelengths at increments of 6 um across a 36 μm range. Maximum intensity projections were created for each channel, and binary masks were created for endothelial cell and neutrophil signal using global thresholding and a custom MATLAB script (MathWorks, 2024b). This script counted neutrophils adherent to the endothelial cell surface, and calculated adhesion fraction by dividing the neutrophil count after washing by the pre-wash neutrophil count. Five technical replicates were performed per condition, across three independent experiments per cell type.

#### Partial Least Squares Regression

A three-component PLSR model was created to fit protein secretion data as a dependent variable to the independent variable of neutrophil adhesion [9]. MATLAB software (MathWorks, 2024b) was used to create and validate this model. R^2^Y captures the percent variance explained by the model in fitting to neutrophil adhesion, we chose the number of components *N* to include in our model such that:

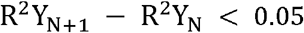

(For our model: Three components, R^2^Y_3_ = 0.718 vs four components, R^2^Y_4_ = 0.758)

### Statistical Analysis

For all experiments, data were pooled from at least three independent biological replicates. Median absolute deviation (MAD) outlier testing was performed in MATLAB on protein secretion data, values >3 MADs from the median were removed and replaced with linear interpolation. One-way ANOVA with Tukey-Kramer post-hoc tests was conducted using MATLAB software. Two-sided Welch’s t-tests were performed using Excel software. P-values are labeled as *p<0.05 unless otherwise noted.

## Results

### 3.1 Endothelial cell source impacted secretion of inflammatory and injury-associated proteins

Endothelial cells at rest are responsible for maintaining an anti-inflammatory state by preventing leukocyte recruitment and coagulation [10]. We sought to determine if endothelial cells from different vascular bed sources exhibited varying basal secretion of inflammatory proteins. For this study we selected a panel of four endothelial cell types: human umbilical artery (HUAEC), human umbilical vein (HUVEC), human dermal microvascular (HDMEC), and human pulmonary microvascular (HPMEC). Each cell type was cultured in a 96-well plate to form a confluent monolayer. We collected supernatant after 16 hours and used multiplexed ELISA to measure basal secretion for a panel of 19 inflammatory and injury markers. Endothelial cells displayed pronounced vascular-bed specific differences in basal secretion of inflammatory and injury-associated proteins (Fig 1). Among the four endothelial sources examined, umbilical vein endothelial cells consistently exhibited the highest basal secretion across most analytes, whereas pulmonary microvascular endothelial cells showed comparatively low overall secretion. Several proteins, including IFN-γ, IL-12p70, syndecan, angiopoietin-1, and IL-10, were below the limit of detection in pulmonary microvascular endothelial cells. Arterial and dermal endothelial cells exhibited similar secretion profiles, differing significantly only in MIP-1α expression (Supplementary Figure 1). Differences in secretion were not restricted to specific protein function (pro-inflammatory, anti-inflammatory, or adhesion), suggesting that endothelial source influenced overall secretory activity rather than selectively altering inflammatory or adhesion-related pathways. These results indicate that endothelial cell sources demonstrate substantial differences in basal protein secretion for immune and injury relevant factors, suggesting heterogeneous inflammatory signaling across vascular beds.

**Figure 1:**
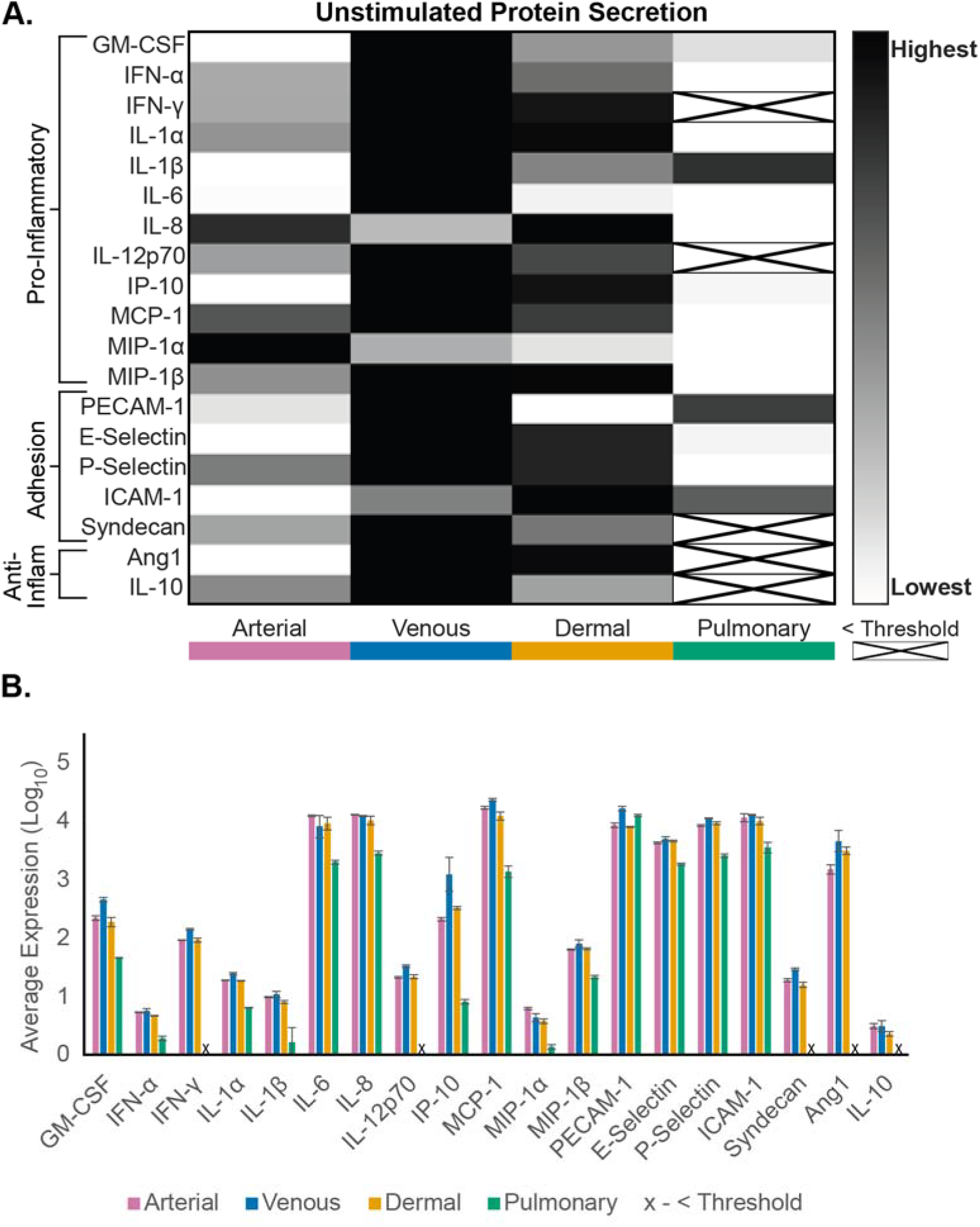
Unstimulated endothelial cells exhibit heterogeneous inflammatory protein secretion. Multiplexed ELISA of conditioned media from unstimulated umbilical artery (HUAEC), umbilical vein (HUVEC), dermal microvascular (HDMEC), and pulmonary microvascular (HPMEC) endothelial cells. Conditioned media was collected from 1 well per cell type after 16 hours incubation. Samples were analyzed with a Luminex Magpix device and a custom ProcartaPlex inflammation panel. Data are represented as **(A)** Heatmap normalized across the range for each protein, with darker tones corresponding to higher expression and less expression for lighter tones. Factors that measured below the limit of detection are indicated with an X. **(B)** log_10_(average expression) ± SEM for N=4 independent experiments. Factors that measured below the limit of detection are indicated with an X. Full statistical analysis with one-way ANOVA and Tukey-Kramer pairwise comparisons testing was done for average expression across all cell types, with results presented in Supplemental Figure 1.

### 3.2 Endothelial cells differentially upregulate inflammatory factor secretion in response to TNF activation

TNF is a known inflammatory activator of the endothelium and causes upregulation of cytokines and adhesion proteins. We wanted to characterize inflammatory responses to TNF-activated endothelial cells and determine how vascular origin impacts secretion profiles. Endothelial cells were cultured as described previously and activated with TNF for 16 hours before supernatant collection and measurement with multiplexed ELISA (Fig 2A). Although venous endothelial cells maintained the highest absolute secretion for many proteins following stimulation, pulmonary microvascular endothelial cells exhibited the largest increase in secretion upon stimulation compared to baseline. Six factors were identified as having the largest relative increases in expression: GM-CSF, IL-1β, IL-6, IP-10, E-selectin, and ICAM-1 (Fig 2B). Pulmonary cells showed the largest increases of all cell types for all factors except E-selectin. Comparatively, umbilical artery cells exhibited the smallest fold-change differences of all cell types; most factors were increased less than two-fold. Notably, dermal microvascular endothelial cells diverged substantially from arterial endothelial cells despite their similar unstimulated secretion profiles, displaying greater upregulation of GM-CSF, E-selectin, and ICAM-1 following TNF exposure. These findings demonstrate that basal endothelial phenotype does not necessarily predict inflammatory responsiveness and highlight pulmonary endothelial cells as especially responsive during cytokine-mediated activation.

**Figure 2:**
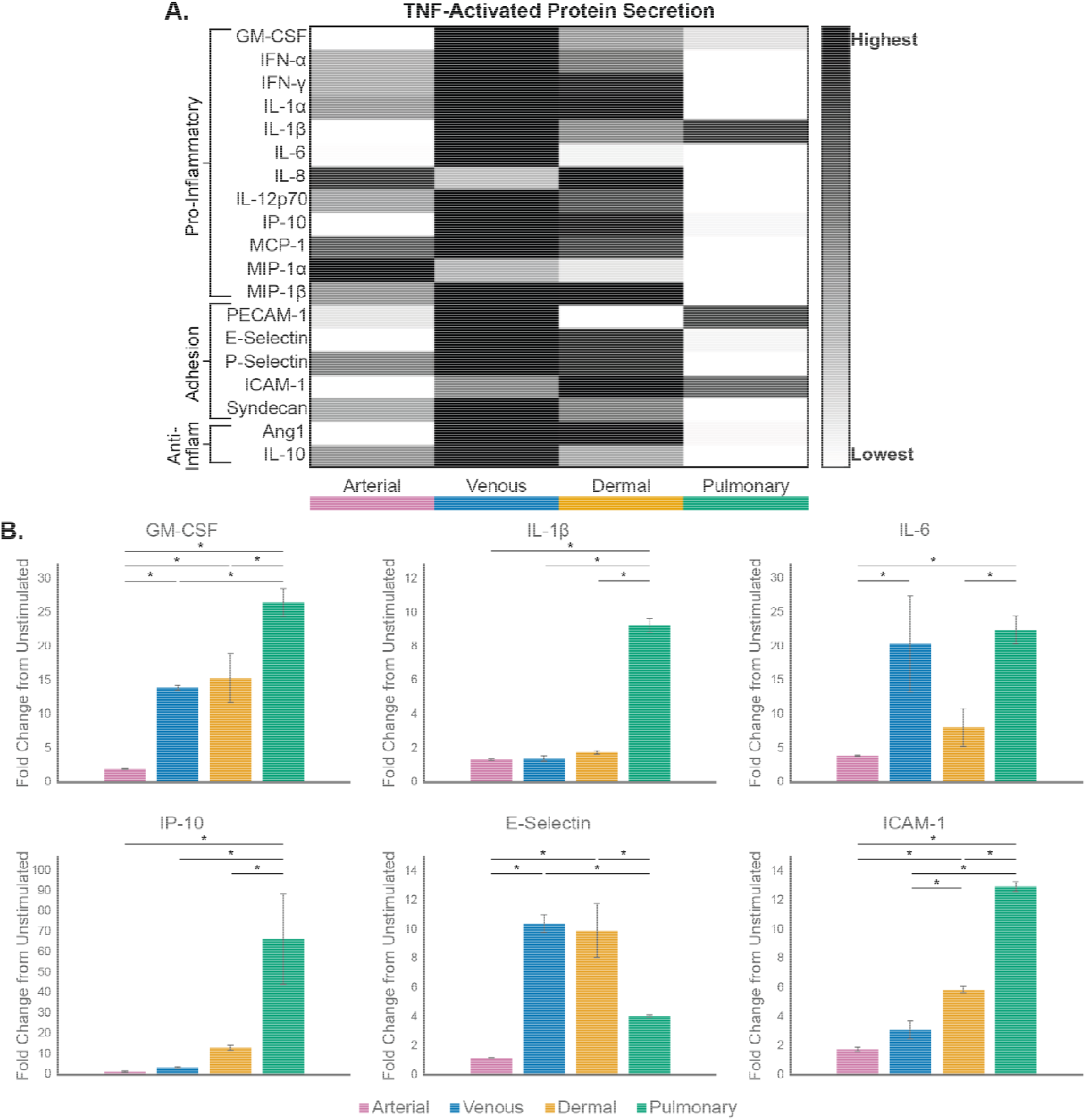
TNF-activated endothelial cells exhibit heterogeneous inflammatory protein secretion. Multiplexed ELISA of conditioned media from TNF-stimulated (50ng/mL TNF in media) umbilical artery (HUAEC), umbilical vein (HUVEC), dermal microvascular (HDMEC), and pulmonary microvascular (HPMEC) endothelial cells. Conditioned media was collected from 1 well per cell type after 16 hours incubation. Samples were analyzed with a Luminex Magpix device and a custom ProcartaPlex inflammation panel. **(A)** Heatmap normalized across the range for each protein, with darker tones corresponding to higher expression and less expression for lighter tones. **(B)** Fold-change expression differences from unstimulated control for factors with >8 fold increase by at least one cell type. Data are average fold change-expression for N=4 independent experiments. One-way ANOVA with Tukey-Kramer pairwise comparisons testing was done for each condition, with asterisks indicating statistical significance between conditions (*p<0.05).

### 3.3 P. aeruginosa inflammatory responses are dependent on endothelial cell source

With our findings of heterogeneous responses to a simple cytokine-mediated inflammatory stimulus, we next sought to characterize if these differences extended to a more complex bacterial infection. We repeated our multiplexed ELISA analysis on endothelial cells that were exposed to *Pseudomonas aeruginosa* for 16 hours (Fig 3A). Umbilical vein cells retained the highest absolute expression of most factors, and pulmonary endothelial cells demonstrated the lowest secretion overall secretion for most factors. As seen with TNF stimulation, GM-CSF, IL-1β, IL-6, IP-10, E-selectin, and ICAM-1 had the largest relative increases from basal expression and wide variation across cell types in response to bacterial activation (Fig 3B). Pulmonary microvascular endothelial cells showed the largest increases in protein secretion, with significantly greater fold-change expression than all other endothelial cell sources for GM-CSF, IL-1β, IP-10, and ICAM-1. Interestingly, dermal microvasculature had significantly greater expression of IL-6 than pulmonary endothelial cells following *P. aeruginosa* activation, opposite the result following TNF activation. These findings reinforce that endothelial cell origin drives distinct inflammatory responses and suggest that different vascular beds exhibit varying responses depending on stimulus type.

**Figure 3:**
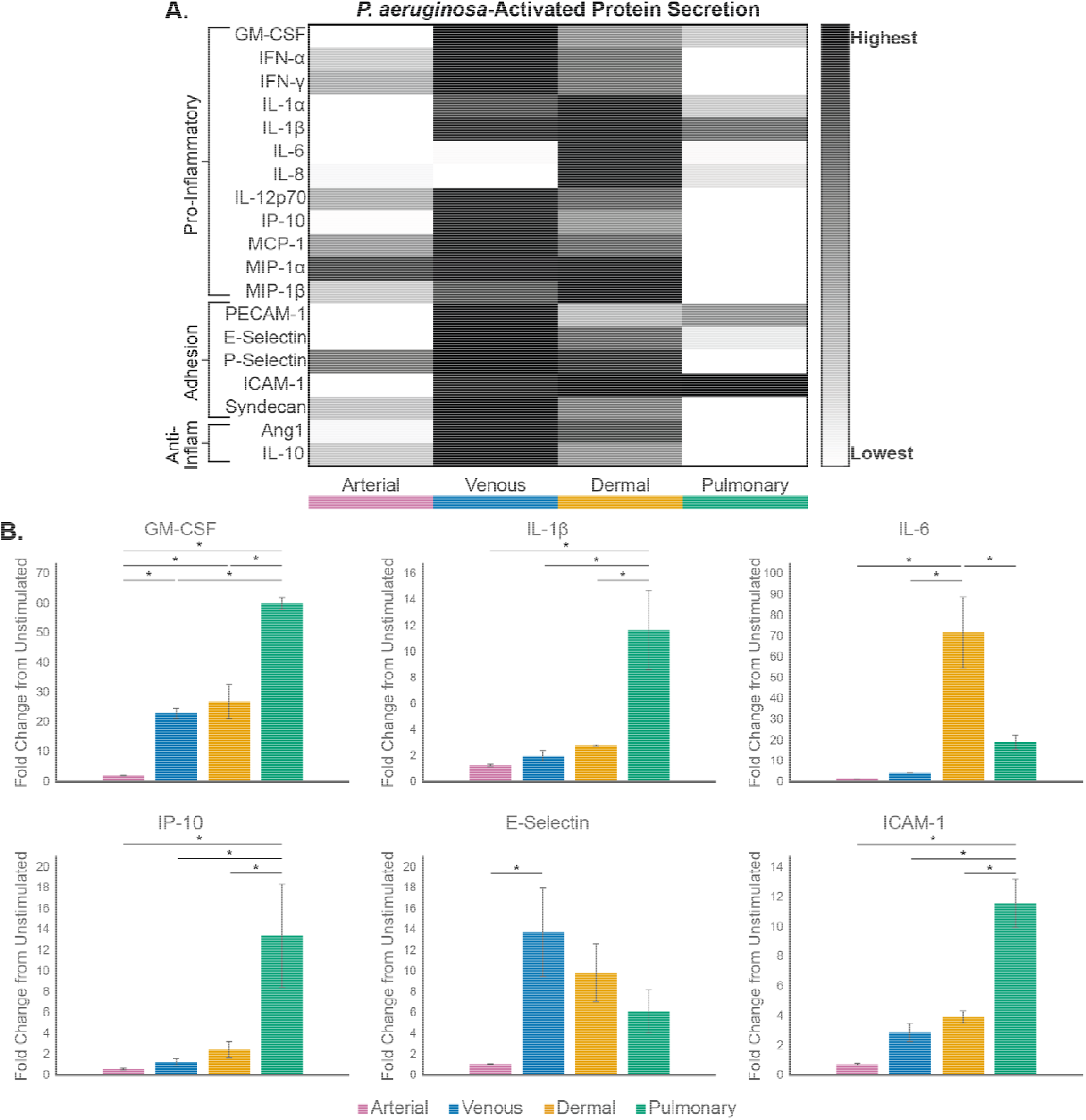
*P. aeruginosa*-activated endothelial cells exhibit heterogeneous inflammatory protein secretion. Multiplexed ELISA of conditioned media from *P. aeuriginosa*-stimulated (0.05 OD in media) umbilical artery (HUAEC), umbilical vein (HUVEC), dermal microvascular (HDMEC), and pulmonary microvascular (HPMEC) endothelial cells. Conditioned media was collected from 1 well per cell type after 16 hours incubation. Samples were analyzed with a Luminex Magpix device and a custom ProcartaPlex inflammation panel. **(A)** Heatmap normalized across the range for each protein, with darker tones corresponding to higher expression and less expression for lighter tones. **(B)** Fold-change expression differences from unstimulated control for factors with >8 fold increase by at least one cell type. Data are average fold change-expression for N=4 independent experiments. One-way ANOVA with Tukey-Kramer pairwise comparisons testing was done for each condition, with asterisks indicating statistical significance between conditions (*p<0.05).

### 3.4 Endothelial cells exhibit distinct responses to TNF and P. aeruginosa activation

*P. aeruginosa* induced a distinct inflammatory profile that partially overlapped with TNF-mediated activation, with some vascular-bed specific differences depending on the stimulus provided (Fig 4). IL-1β and E-selectin were largely indifferent to stimulus type, and the differences between endothelial cell sources were relatively consistent across activations. In contrast, GM-CSF, IL-6, IP-10, and ICAM-1 displayed strong stimulus dependence. TNF preferentially induced higher IP-10 and ICAM-1 expression, while *P. aeruginosa* more strongly stimulated GM-CSF production. IL-6 responses were particularly vascular-bed specific: arterial, venous, and pulmonary endothelial cells responded more strongly to TNF, while dermal microvascular endothelial cells preferentially upregulated IL-6 following bacterial exposure. Together, these data indicate that endothelial inflammatory heterogeneity is dependent not only on vascular origin, but also from stimulus-specific activation.

**Figure 4:**
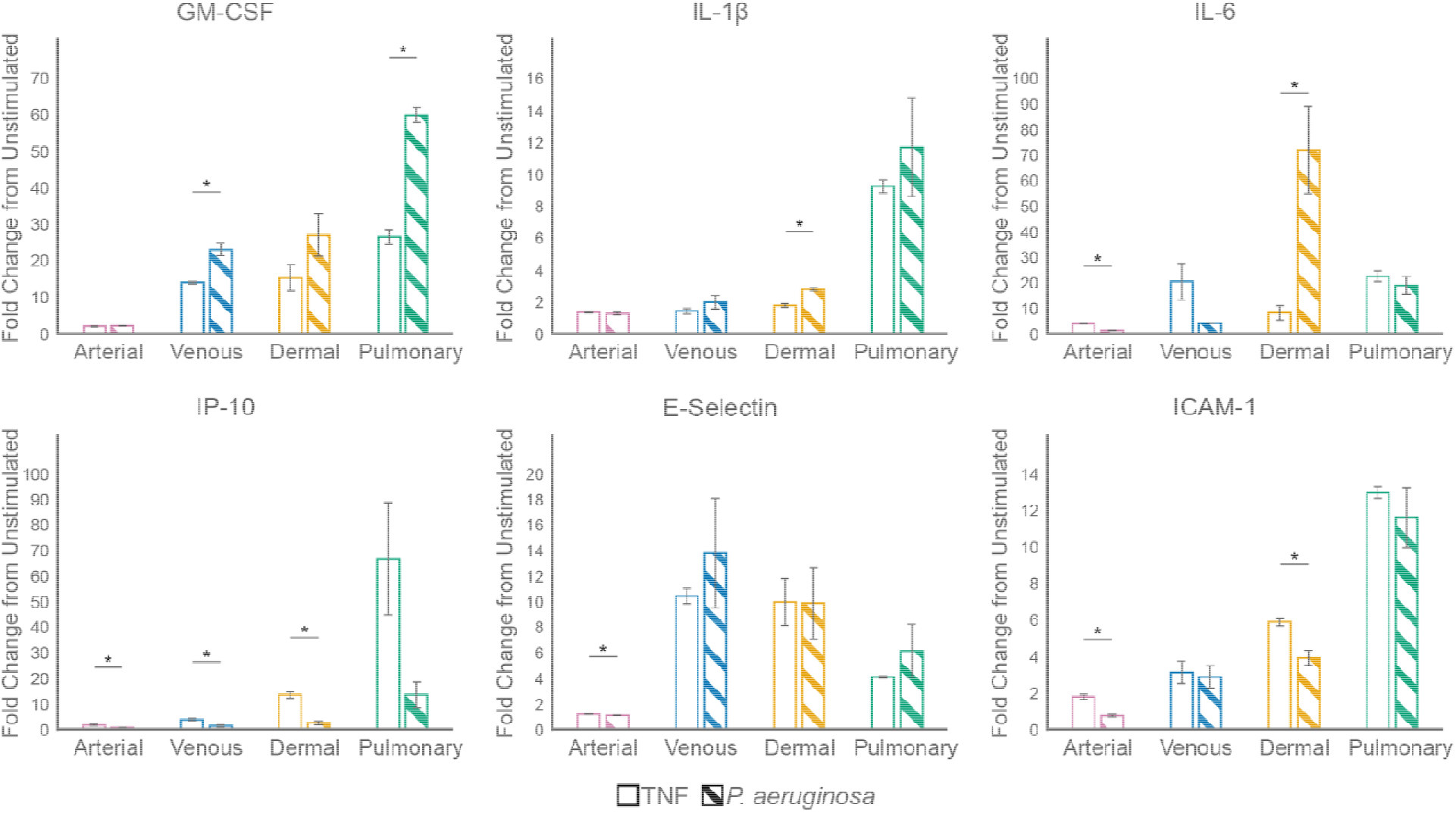
TNF and *P. aeruginosa* activation differentially altered endothelial cell inflammatory secretion by cell type. Data represent fold-change expression differences in protein secretion from Figure 2 and Figure 3, plotted to compare TNF and *P. aeruginosa* specific differences. Welch’s t-tests were performed comparing each activation, with asterisks indicating statistical significance between activations for a given cell type (*p<0.05). Error bars indicate average ± SEM.

### 3.5 Endothelial cell source affects neutrophil adhesion

Having identified unique inflammatory profiles for various endothelial cells, we wanted to determine if those differences would meaningfully contribute to changes in downstream immune cell recruitment. Neutrophils adhering to endothelium is a critical first step in the neutrophil adhesion cascade, and we therefore wanted to determine if the endothelial heterogeneity extended to differences in neutrophil adhesion to the endothelial cell surface. Endothelial cell monolayers were activated with either TNF or *P. aeruginosa*, and primary neutrophils were added to adhere. Non-adherent neutrophils were removed with successive wash steps (Fig 5A). Neutrophil adhesion was quantified as the fraction of adherent neutrophils from the base loading density (Fig 5B). Umbilical vein endothelial cells exhibited significantly higher adhesion in the control condition than umbilical artery or dermal microvascular endothelial cells (Fig 5B). TNF and *P. aeruginosa* activation generally resulted in only modest increases in neutrophil adhesion and reached statistical significance only in TNF-treated venous endothelial cells (Fig 5C). Collectively, these findings suggest that endothelial source contributes more strongly to baseline neutrophil recruitment capacity than to adhesion to inflamed endothelium under the conditions tested.

**Figure 5:**
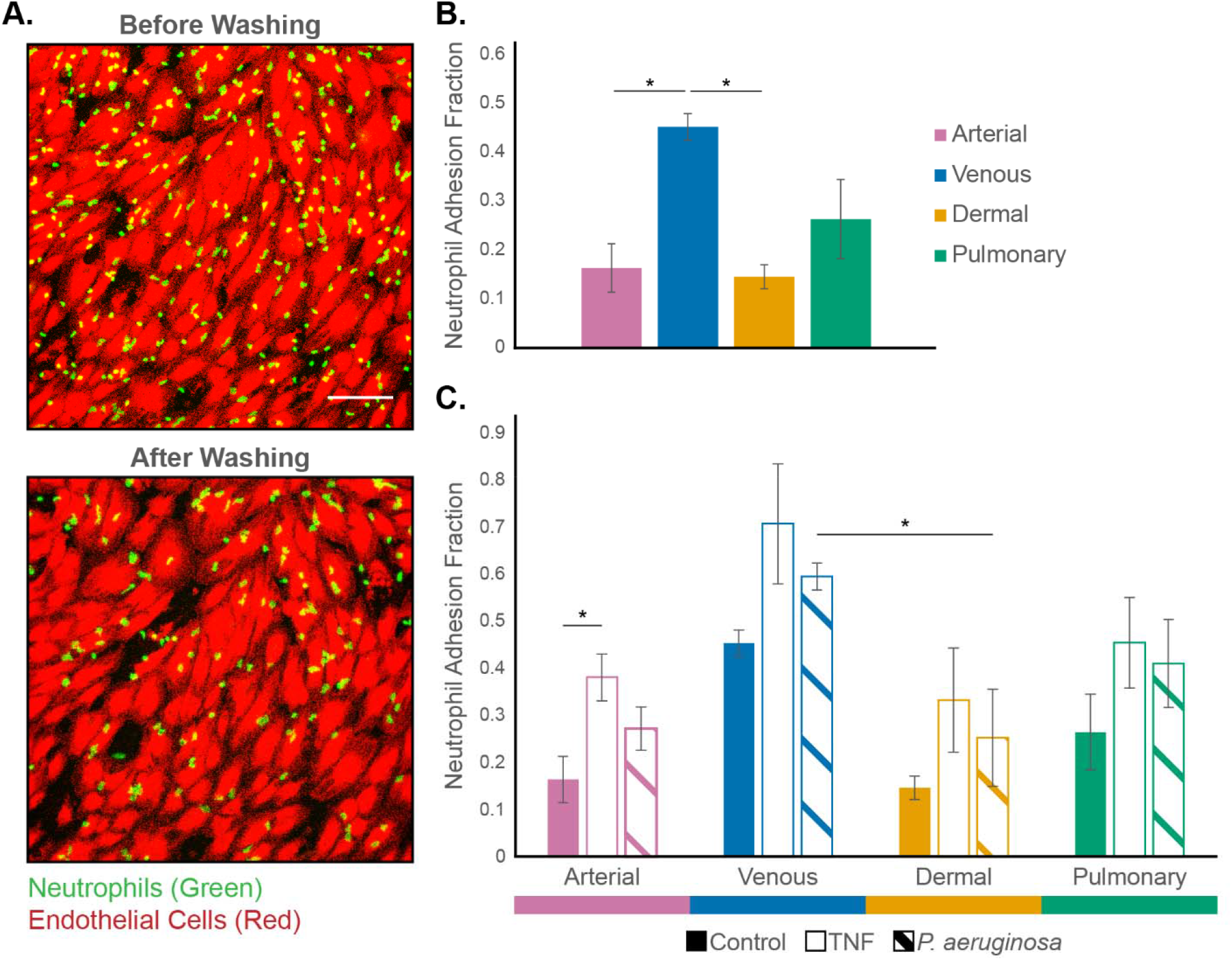
Neutrophil adhesion differs across endothelial cell sources. Endothelial cells were cultured in 96-well plates and activated with control, TNF(50 ng/mL) or *P. aeruginosa* (0.05 OD) media for 2 hours. Neutrophils isolated from whole blood were added to each well and incubated for 15 minutes, then imaged before and after 5x PBS washes. **(A)** Representative images of endothelial cells (red) and neutrophils (green) before and after washing to remove non-adherent cells. (Scale bar = 100 μm) **(B, C)** Neutrophil adhesion fraction was quantified by counting neutrophils adherent after washing and dividing by the neutrophil count before washing. Data represent average adhesion fraction ± SEM for 3 independent experiments per cell type. One-way ANOVA with Tukey-Kramer pairwise comparisons testing was done for each condition, with asterisks indicating statistical significance between conditions (*p<0.05).

### 3.6 Partial Least Squares Regression (PLSR) reveals secreted factors that correlate with neutrophil adhesion

Finally, we sought to elucidate the relationship between secreted factors and neutrophil adhesion and, therefore, employed a dimension-reduction analysis to identify key correlations between the two. Specifically, we applied partial least squares regression (PLSR) using protein secretion profiles (independent variable, X) to predict neutrophil adhesion outcomes (dependent variable, Y). PLSR combines aspects of principal component analysis and linear regression, where a complex set of highly correlated predictor variables can be simplified to smaller number of components that maximize the fit of X onto Y. These components can then be analyzed to determine which independent variables are the most important to maximizing the model’s fit, this can be quantified through Variable Importance in Projection (VIP) scores. A three-component PLSR model adequately captured the variance in adhesion (R^2^=0.718), without substantial improvements from additional components (Fig 6A, B). VIP analysis revealed 5 factors with a VIP score of near or greater than 1: E-Selectin (2.63), PECAM1 (2.43), ICAM-1(1.82), IL-6 (1.03), and IL-8 (0.97) (Fig 6C). E-selectin, PECAM1, and ICAM-1 are the strongest positive predictors of neutrophil adhesion. IL-6 and IL-8 also contributed substantially to the model, though they exhibited negative correlation with adhesion. Notably, three proteins most strongly associated with adhesion (E-selectin, ICAM-1, and IL-6) were also among the factors exhibiting the largest increases from baseline and showed the greatest endothelial heterogeneity across vascular beds. These findings link endothelial source-dependent inflammatory signaling to functional differences in neutrophil recruitment.

**Figure 6:**
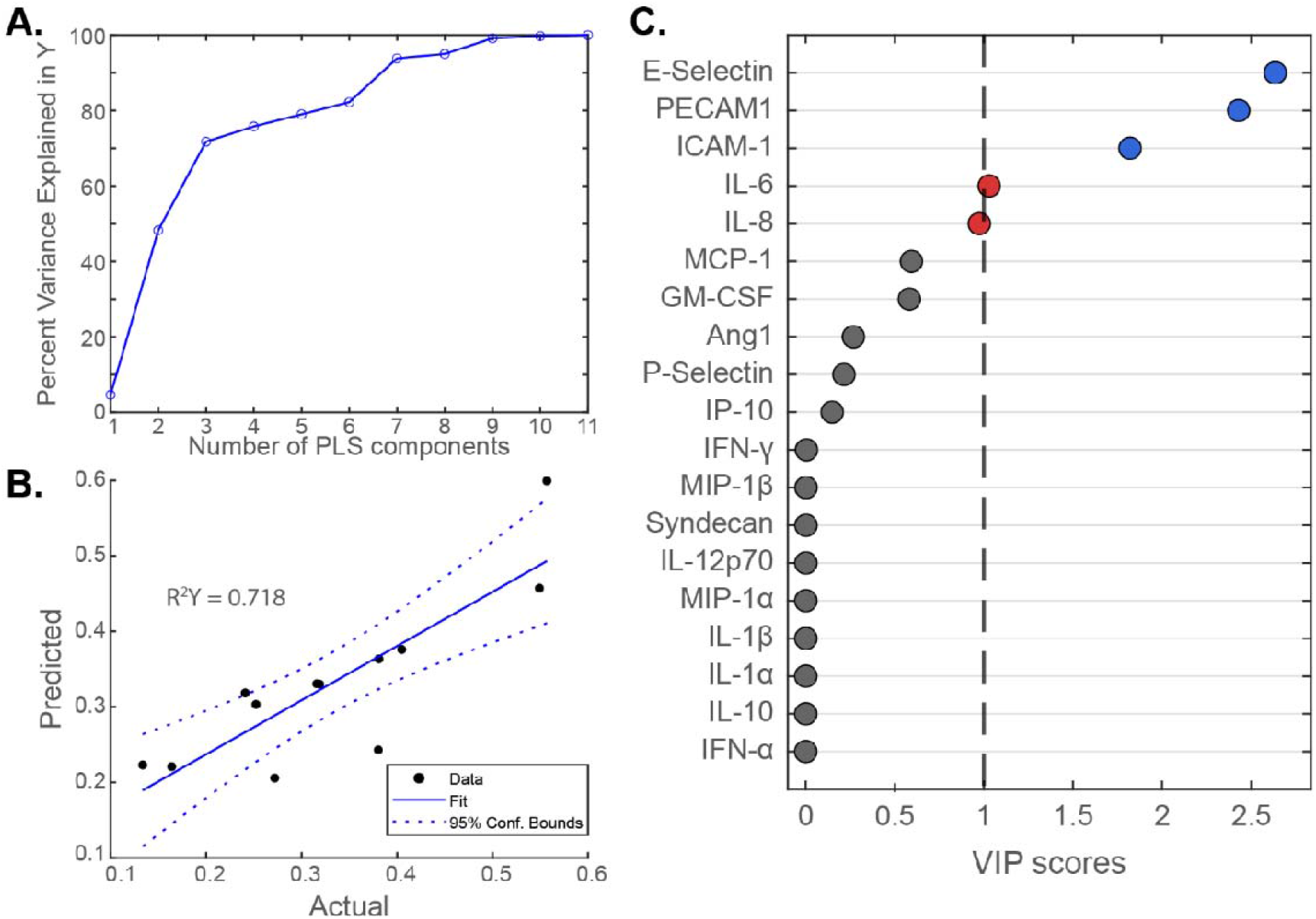
Partial least squares regression identifies secreted factors most associated with neutrophil adhesion changes. Secreted protein data (independent variable, X) was fit to neutrophil adhesion data (dependent variable, Y) **(A)** The number of components to include in the model was determined by calculating the percent variance in Y (neutrophil adhesion) for N=1-11 components, and finding the point where adding an additional component improved the variance explained by <5% (N=3). **(B)** Diagnostic plot for actual vs fitted values for the 3-component model **(C)** VIP scores of each secreted factor ranked highest to lowest, with VIP scores near or greater than 1 filled in to indicate the direction of correlation (blue=positive, red = negative). Factors significantly below VIP=1 are filled in gray.

## Discussion

In this study, we found that endothelial heterogeneity significantly influences inflammatory signaling and downstream immune interactions. Endothelial cells (ECs) derived from different vascular beds displayed distinct basal secretion profiles and varied sensitivity to inflammatory stimuli. Notably, venous endothelial cells exhibited the highest secretion of most inflammatory markers, and pulmonary cells showed low basal secretion with the largest relative increases following activation. Furthermore, different inflammatory stimuli of TNF and *P. aeruginosa* produced overlapping but distinct secretion patterns, particularly for factors GM-CSF, IL-1β, IL-6, IP-10, E-selectin, and ICAM-1. We also saw endothelial cell source impacted baseline neutrophil adhesion, with venous endothelial cells having significantly higher adhesion than other cell types. Finally, we used a regression model to map protein-secretion data to neutrophil adhesion to identify the factors that best predicted neutrophil recruitment trends: E-selectin, ICAM-1, PECAM1, IL-6, and IL-8. Collectively, these findings suggest that endothelial source has a large impact on vascular inflammatory responses and downstream immune cell interaction. These observations support an emerging view of endothelial cells as active regulators of immune responses whose behavior is strongly influenced by vascular-bed specialization rather than uniformly conserved across tissues.

Previous studies have revealed extensive endothelial diversity across tissues using single-cell analysis, including distinct arterial, venous, and tissue-specific endothelial subpopulations with specialized molecular signatures and signaling functions [4,6,7,11]. The lung microvasculature in particular possesses unique functional demands because excessive inflammatory activation can impair gas exchange and tissue function, but dynamic immune responses are necessary as the lungs face constant risk of environmental and pathogen exposure[12,13]. Our study is consistent with this framework, as we observed pulmonary microvasculature had low unstimulated secretion with the most dynamic response to inflammatory stimulus. Previous work has shown umbilical artery endothelial cells express less adhesion molecules and lower monocyte recruitment following TNF activation than umbilical vein endothelial cells [14]. We saw similar trends in this study, with arterial cells showing the lowest relative increases in protein secretion for most factors and venous endothelial cells having the highest expression for most factors measured.

While we observed heterogeneous responses to inflammation across endothelial cell sources, we also saw further variation dependent on stimulus type. Interestingly, the same six factors were the most highly upregulated for both TNF and *P. aeruginosa* stimulation: GM-CSF, IL-1β, IL-6, IP-10, E-selectin, and ICAM-1. TNF stimulation preferentially increased IP-10 and ICAM-1 expression, whereas bacterial stimulation more strongly induced GM-CSF production and caused vascular-bed specific differences in IL-6 expression. These findings likely reflect differences in signaling pathways activated by each stimulus. TNF predominantly initiates inflammatory responses through TNF receptor-mediated activation of NF-κB[10], while bacterial exposure activates pathogen-sensing mechanisms through pattern-recognition receptors[15,16]. TNF-mediated NF-κB signaling is associated with endothelial activation and directly leads to increases in ICAM1 and IP-10, corresponding with our results [17,18]. Bacteria responses are more complicated and robust, because multiple toll-like receptors (TLRs) can be activated leading to varied responses, such as *P. aeruginosa* activating TLR2, TLR4, and TLR5 [19,20]. TLR activation can lead to increasing GM-CSF expression and a positive feedback loop of increased TLR expression and sensitivity to bacterial activation [21]. Recent work has demonstrated endothelial cells adopt distinct inflammatory states to different cytokine stimuli, producing different transcription and secretion profiles rather than one uniform activated state[22]. Lastly, we saw that despite similar unstimulated secretion of most factors, arterial and dermal microvasculature endothelial cells diverged in their responses to inflammatory stimuli. Overall, these observations indicate that a combination of intrinsic vascular identity and external inflammatory cues dictate vascular activation.

Our PLSR model identified three adhesion molecules to be most strongly correlated with neutrophil adhesion: ICAM-1, E-selectin, and PECAM1. Each of these is well-established in neutrophil-endothelial cell interactions, with E-selectin driving neutrophil rolling, ICAM-1 causing firm adhesion, and PECAM1 contributing to diapedesis[2]. In contrast, previous work has shown secreted adhesion molecules can interfere with leukocyte adhesion through competitive inhibition, though these studies used exogenous protein levels higher than physiologically relevant plasma concentrations [23,24]. Therefore, we theorize that increased secretion of adhesion molecules corresponded to an overall increased state of endothelial activation that overcame the contributions of this competitive inhibition. Future work will be necessary to confirm endothelial heterogeneity of membrane-expression for these factors, and corresponding impacts to neutrophil behavior. Our analysis also showed that increased IL-6 and IL-8 expressions are negatively correlated with neutrophil adhesion. Both cytokines are typically associated with pro-inflammatory effects, but there is evidence suggesting their impact in neutrophil-endothelial cell interactions is not straightforward. We have previously demonstrated a complex interaction between endothelial cells, neutrophils, and IL-6 in response to infection where blocking IL-6 prevents all neutrophil extravasation from a model blood vessel[25] but high levels of exogenous IL-6 have deleterious effects on neutrophil response and are associated with decreased ICAM-1 [26]. IL-6 has been found to exert both pro- and anti-inflammatory effects based on concentration and context [27]. Interestingly, shortly after IL-8 was first discovered it was noted to have anti-inflammatory effects and cause clearance of neutrophils from the endothelial surface[28], though now it is mostly recognized to increase neutrophil recruitment to the endothelial surface [29,30]. This apparent discrepancy may be explained by how IL-8 is presented to neutrophils *in vivo*, where glycosaminoglycans capture IL-8 and present it to rolling neutrophils to prime them for firm adhesion and transmigration [30]. Soluble IL-8 binding to neutrophils can interfere with stable adhesion as they quickly respond and change receptor behavior before they are in proximity to endothelial cells, and IL-8 exposure has been shown to desensitize neutrophils. Of the factors our PLSR model identified as most predictive of neutrophil adhesion, E-selectin, ICAM-1, and IL-6 were also found to be among the factors most impacted by endothelial cell origin during an inflammatory response. This further reinforces the impact endothelial cell heterogeneity can have on downstream neutrophil interactions.

In this study, we demonstrate the impact that endothelial cell origin can have on inflammatory responses and downstream neutrophil interactions. To our knowledge, this is the first study to analyze the variation of inflammatory secretomes in the context of endothelial heterogeneity and how those differences contribute to neutrophil recruitment changes. Our study employed a simple *in vitro* methodology to model acute inflammation, but future work is necessary to confirm and characterize these differences in more complex models that incorporate vessel geometry, extracellular matrix, and physiological fluid flow, factors known to influence endothelial cell activation and behavior. Our study is also constrained by a limited number of endothelial cell sources, so donor-to-donor heterogeneity likely impacted our findings, and further work will be necessary to pinpoint specific vascular-bed differences. Despite these limitations, our work highlights endothelial cell selection as a critical design consideration when developing immune *in vitro* models. Future research may reveal further downstream impacts of vascular heterogeneity on endothelial cell-coordinated immunity, including neutrophil effector functions and other immune cell interactions.

