## Supplemental Data for "Endothelial Heterogeneity Across Vascular Beds Impacts Inflammatory Signaling and Neutrophil Adhesion"

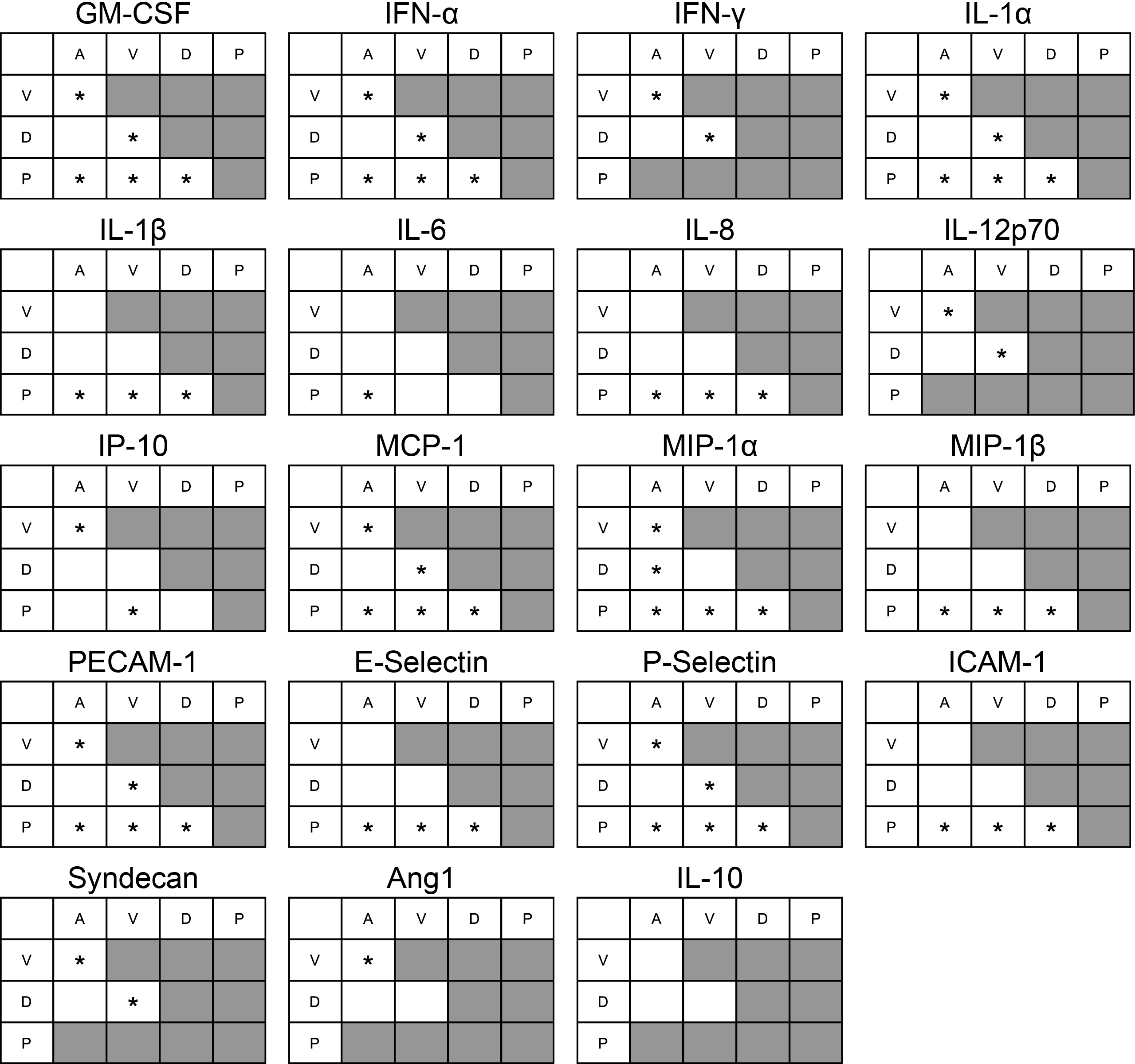
 **Supplemental Figure 1: Full statistical analysis for secretion data of unstimulated endothelial cells.** One-way ANOVA and Tukey-Kramer post hoc tests were conducted for unstimulated expression of the 19 factors measured by multiplexed ELISA. Pairwise comparisons between cell types for all factors are summarized in the table (A-umbilical artery, V-umbilical vein, D-dermal microvasculature, P-pulmonary microvasculature), where asterisks denote significantly different expression levels between cell types (*<0.05). Pulmonary expression was below the limit of detection for 5 factors, and these comparisons were excluded from the analysis: IFN-γ, IL-12p70, syndecan, angiopoietin-1, and IL-10.
